# Chromosome-level genome assembly and annotation of the parthenogenetic nematode *Acrobeloides nanus*

**DOI:** 10.64898/2026.03.06.710095

**Authors:** Nadège Guiglielmoni, Laura I. Villegas, Michael Paulini, Lewis Stevens, Anja Schuster, Christian Becker, Kerstin Becker, Mark Blaxter, Philipp H. Schiffer

**Affiliations:** worm~lab, Institut für Zoologie, Universität zu Köln, 50674 Cologne, Germany; Tree of Life Programme, Wellcome Sanger Institute, Wellcome Genome Campus, Cambridge CB10 1SA; Biological and Medical Research Center (BMFZ), Medical Faculty and University Hospital Düsseldorf, Heinrich Heine University Düsseldorf, Düsseldorf, Germany; West German Genome Center (WGGC), Medical Faculty and University Hospital Düsseldorf, Heinrich Heine University Düsseldorf, Düsseldorf, Germany; Cologne Center for Genomics (CCG), Medical Faculty, University of Cologne, Cologne, Germany; West German Genome Center (WGGC), University of Cologne, Cologne, Germany; Biodiversity Genomics Center Cologne (BioC^2^), Cologne, Germany

## Abstract

Acrobeloides nanus is a species of widely distributed bacteria-feeding nematodes belonging to the family Cephalobidae. It reproduces parthenogenetically, making it a valuable system for studying the consequences of obligate asexuality, and exhibits early developmental processes that differ significantly from the model organism Caenorhabditis elegans. Additionally, the species has demonstrated remarkable resistance in extreme conditions, surviving near-complete desiccation through anhydrobiosis.

This new chromosome-level genome assembly provides a high-quality reference to study these traits. Combining Nanopore, amplified PacBio HiFi, and Hi-C sequencing, we generated a chromosome-level assembly spanning 188.9 Mb over six chromosomes. The genome features a 48.1% repeat content, dominated by Mutator-like elements, and contains 24,912 predicted genes. Structural analysis reveals the conservation of ancestral nematode linkage groups. This resource significantly improves upon previous fragmented assemblies, offering a new framework for identifying genomic adaptations in the family Cephalobidae.

## Background

*Acrobeloides* is a genus of free-living nematodes classified within the infraorder Tylenchina (order Rhabditida, Clade IV) [1, 2, 3], belonging to the family Cephalobidae [4]. This group is ubiquitous, classified as bacteria-feeding nematodes [5], and is one of the more abundant nematode groups across diverse terrestrial environments, including fields, forests, and arid environments [5, 6, 7, 8]. The genus *Acrobeloides* contains both sexually reproducing and parthenogenetic species, making it a valuable system for studying the evolutionary transition to asexuality from a developmental [9] and ecological perspective [10].

*Acrobeloides* species are known to be particularly resilient to adverse conditions, such as drought, often surviving by entering a dormant state [11]. Specifically, anhydrobiotic abilities were documented in the parthenogenetic species *Acrobeloides nanus*, capable of surviving near-complete desiccation by adopting a temporary coiled structure [12, 13, 14]. *A. nanus* has been used to measure responses to abiotic stress induced by varying copper and pH levels [10, 15]. Partly based on these studies it was suggested that *A. nanus* would be useful for toxicity assessments under natural conditions [16].

From an evolutionary developmental point of view *A. nanus* has been an intriguing system, as a regulative potential in embryonic development in Nematoda was first described in this species [17]. Its early development on the cellular level is thus significantly different from the nematode model system *Caenorhabditis elegans* [18, 19], allowing for studies into nematode evolution of development [20]. Cephalobidae have elaborate head morphologies [21], which are thought to be ecological adaptations to different bacterial food sources [22] and fascinating from a standpoint of post-embryonic development. As the exact relationships within the family are unresolved, with the possibility that the genera *Acrobeloides* and *Cephalobus* should be united into one genus (see e.g. [23]), comparative studies into ecology, evolution and development are currently impeded by phylogenetic uncertainty.

Previous genomic studies of *A. nanus* have been limited by the quality of available resources; two highly fragmented short-read assemblies were previously released, offering only kilobase-level contiguity [20, 24]. These fragmented assemblies have restricted comprehensive analyses of chromosome evolution, repetitive elements, as well as detailed studies in ecology, evolution and development.

Here we present a high-quality, chromosome-level collapsed haploid assembly of *Acrobeloides nanus*. This resource provides the necessary scaffold to enable detailed comparative studies addressing genome evolution in the context of parthenogenesis and anhydrobiosis, resolve taxonomic questions and offers a new framework for identifying genomic adaptation for this nematode’s environmental resilience.

## Methods

### Nanopore R10.4 sequencing

High-molecular-weight DNA was extracted as described in [25]. Briefly, worms were pelleted from agar plates, decontamined with sucrose and dissolved in a cetyltrimethylammonium bromide (CTAB) buffer (polyvinylpyrrolidone 2%, Tris-HCl 100 mM, ethylenediaminetetraacetic acid 25 mM, NaCl 2 M, CTAB 2%) supplemented with 25 *µ*L of proteinase K (Zymo Research D3001-2). Samples were incubated for 1 hour at 60°C, and then for another 10 minutes with 1.0 M potassium acetate. DNA was purified with phenol-chloroform-isoamyl alcohol 25:24:1, chloroform-isoamyl alcohol 24:1 and AMPure XP beads (Agencourt), and RNA was removed with RNAse cock-tail enzyme mix (Thermo Fischer, AM2286) for 1 hour at 37° C. DNA was fragmented in a 2 mL low-bind round bottom Eppendorf tube using a sterile 3 mm borosilicate bead (Z143928-1EA Merck) by vortexing for 1 minute at maximum speed as described in [26] and short fragments were removed using the Short Reads Eliminator (SRE) (Circulomics, Pacific Biosciences). DNA concentrations were quantified using a Qubit 4 fluoremeter with 1X dsDNA kit. Nanopore libraries were prepared using the Ligation Sequencing Kit LSK114 (Oxford Nanopore Technologies) and sequenced on R10.4 MinION flowcells. Basecalling was performed using Dorado v0.3.1 [27] in duplex mode with model dna r10.4.1 e8.2 400bps supv4.2.0 and the reads were converted to fastq using SAMtools v1.6 [28] with the module samtools fastq.

### Ultra-low input amplified Pacific Biosciences HiFi sequencing

Up to 10 individuals were lysed with salt-based extraction buffer (Tris-HCl 100 mM, ethylenediaminetetraacetic acid 50 mM, NaCl 0.5 M and sodium dodecylsulfate 1%) and 5 *µ*L of proteinase K (Zymo Research D3001-2) overnight at 50°C, as described in [25]. DNA was precipitated using NaCl 5 M, yeast tRNA and isopropanol, and incubated at room temperature for 30 minutes, then pelleted at 18,000 g for 20 minutes (4°C). The DNA was washed twice with 80% ethanol and spinned at 18,000 g for 10 min (4°C). The DNA pellet was eluted in elution buffer (D3004-4-10 Zymo Research) and incubated at 50°C for 10 minutes. RNA was removed by incubating with RNAse (Qiagen, 19101) for 1 hour at (37°C). DNA concentrations were quantified using a Qubit 4 fluoremeter with 1X dsDNA kit. The amplified PacBio HiFi library was prepared with the Express 2.0 Template kit and the SMRTbell prep kit 3.0 (Pacific Biosciences, Menlo Park, CA, USA), and sequenced on a Sequel II instrument with 30 hours movie time. PacBio HiFi reads were generated using SMRT Link (v11.1, Pacific Biosciences, Menlo Park, CA, USA) with default parameters.

### Hi-C sequencing

Worms were pelleted and crosslinked in 3% formaldehyde for 45 minutes and quenched in 250 mM glycine for 25 minutes. They were then processed with Arima Hi-C+ kit starting from lysis. The library was sequenced on an Illumina NovaSeq 6000 at the Cologne Center for Genomics.

### Genome assembly

Amplified PacBio HiFi and non-amplified Nanopore reads were assembled using hifiasm v0.19.4 [29] with parameters `-l 3 --ul`. The alignment from BLAST v2.13 [30], the output from BUSCO v5.4.7 [31] against Metazoa odb10, and the sorted mapping of PacBio HiFi reads were provided as input to BlobToolKit v4.1.5 [32] to identify and remove contaminants. PacBio HiFi reads were mapped using minimap2 v2.24 [33] with parameter `-x map-hifi` to subsequently remove alternative haplotypes using purge dups v1.2.5 [34]. Adapters were searched for and removed using the NCBI Foreign Contamination Screen (FCS) with the module FCS-adaptor v0.5.4 [35]. Basic assembly statistics were computed using assembly-stats v1.0.1 [36]. BUSCO v5.8.0 [31] was run against Nematoda odb12 with parameter `-m genome`. *k* -mer comparison plots were computed using KAT v2.4.2 [37] with the module `kat comp` and the PacBio HiFi reads. Hi-C contact maps were generated by processing Hi-C reads using hicstuff v3.1.1 [38] as described previously and visualizing the contacts using the module `hicstuff` `view` with the parameter `-b 500`. Orthologs from the Nematoda odb10 lineage were searched for in genome assemblies using BUSCO v4.1.4 [31] and classification to the ancestral rhabditid linkage groups (Nigon elements) were identified using the online platform of vis alg [39]. Chromosome-level assemblies were checked for residual contamination and corrected using BTK [32], and FCS [40]. Manual curation was performed using JBrowse 2 [41], HiGlass [42] and PretextView [43].

### Repeat and gene annotations

The Extensive *De novo* TE Annotator (EDTA) pipeline v2.0.1 [44] was run with parameters `--sensitive 1 --anno 1 --force 1`. The hardmasked assembly was converted into a softmasked assembly. RNA-seq reads were trimmed using TrimGalore v0.6.10 [45] and mapped to the assemblies using hisat2 v2.2.1 [46]. After sorting the mapped reads using SAMTools v1.6 [28], the output was provided as input to BRAKER v3.0.3 [47] with parameters `--gff3 --UTR off`, which combined predictions from Augustus v3.5.0 [48] and GeneMark-ES v4.68 [49]. The longest isoform was selected using Another Gtf/Gff Analysis Toolkit (AGAT) v0.8.0 [50] with the script `agat sp keep longest isoform.pl.` The protein predictions were evaluated using BUSCO v5.8.0 [31] against the Nematoda odb12 lineages with the parameter `-m protein`.

### Mitochondrial genome assemblies and annotations

The module `findMitoReference.py` from MitoHiFi v3.2.1 [51] was run with parameter `--type mitochondrion` to search for a closely related mitochondrial genome to use as a reference and identified the complete mitochondrial genome of *Acrobeloides varius* (MK559448) [52]. MitoHiFi was run with reads as input (`-r`) and parameters `-o 5 --mitos -f MK559448.1.fasta -g MK559448.1.gb`.

### Synteny

Orthologs were identified within the annotated genes by BUSCO v4.1.4 and the Nematoda odb10 lineage. An Oxford dot plot was constructed with a custom script between *Acrobeloides nanus* and *Acrobeloides maximus* (nxAcrMaxi1.1, GCA 964212105.1) based on these orthologs, as in [53].

### Genome description

We generated ultra-low input PacBio HiFi reads from a few individuals with amplification, Nanopore R10.4 reads from a large pool of individuals, and Hi-C from a large pool of individuals. Sequencing resulted in 33.2 Gb of PacBio HiFi reads, 14.6 Gb of Nanopore reads, 126 millions pairs of Hi-C reads. It should be noted for future reuse that the Hi-C library was highly contaminated with bacteria (as food source) and only 17% of the reads mapped to the assembly. *k* -mer analyses were run on the Nanopore reads, as heterozygous and homozygous peaks were not properly identified in the PacBio HiFi reads due to amplification, and the nuclear genome was predicted to be diploid (Figure 1A).

**Figure 1:**
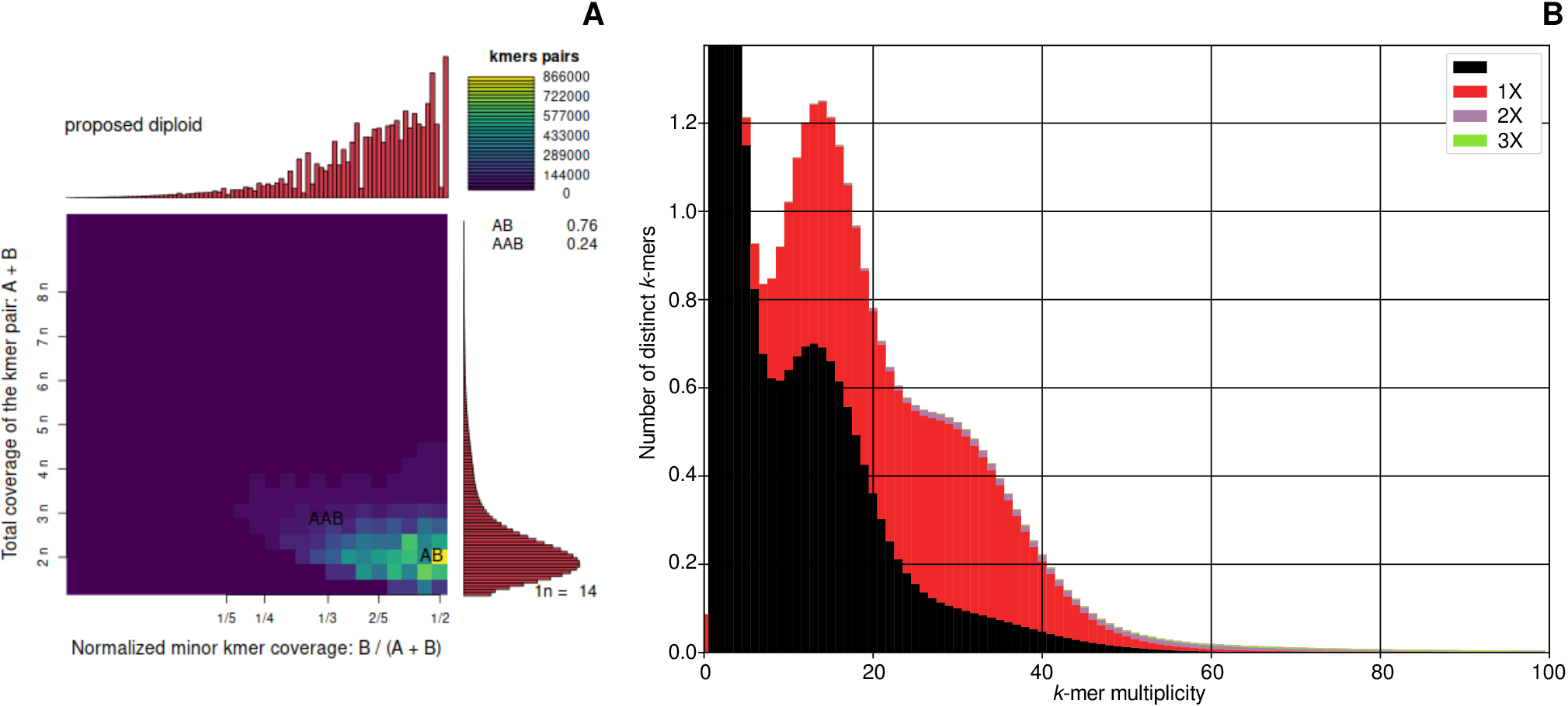
*Acrobeloides nanus* is predicted as diploid. 27-mer analysis of the Nanopore sequencing dataset and the genome assembly. A) Ploidy assessment using Smudgeplot. The strong AB signal at 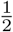 and 2n suggest diploidy. B) Comparison of 27-mers in the Nanopore reads and those integrated in the genome assembly. *k* -mer multiplicity and number of distinct *k* -mers refer to the 27-mers identified in the Nanopore reads. Curves are colored based on the representation of these same 27-mers in the assembly: 0X black (absent from the assembly), 1X red, 2X purple, 3X green.

The nuclear genome assembly of *A. nanus*, available in the European Nucleotide Archive as nxAcrNanu1, was collapsed to represent the two haplotypes into single sequences. The final assembly has homozygous regions included in one copy and heterozygous regions represented in one copy or absent from the assembly, as only one haplotype could be included (Figure 1B). The homozygous peak shows some missing sequences, as 0X; this could be sequences in the Nanopore sequencing dataset and absent from the PacBio HiFi reads, which were used for the initial hifiasm assembly. The initial assembly included a large amount of bacterial contamination, due to the feeding mode of *A. nanus*, which was removed from the final assembly (Figure 2). The final assembly reached a size of 188.9 Mb with 6 chromosome-level scaffolds spanning 98.98% of the assembly (Figure 3A, Table 1). This size is smaller than previous short-read assemblies at 248.1 Mb [20] and 241.6 Mb [24], but the genome size was estimated to 176.6 Mb in [24], close to the size of this long-read assembly. The longer short-read assemblies may be owed to partial haplotype collapsing or contamination. Chromosome candidates have similar sizes ranging from 29.6 to 32.5 Mb. Along with the *k* -mer comparison suggesting a high completeness, the BUSCO score reached 95.6% against the Nematoda odb12 lineage (92.6% single-copy ortholog, 3.0% duplicated orthologs).

**Table 1:** Assembly statistics for *Acrobeloides nanus* nxAcrNanu1. BUSCO orthologs were identified from the Nematoda odb12 lineage.

|  |  |  |
| --- | --- | --- |
| **Assembly size** |  | 188,892,103 bp |
| **Contig number** |  | 101 |
| **Scaffold N50** |  | 31,568,468 bp |
| **Scaffold N90** |  | 29,610,334 bp |
| **Contig N50** |  | 284,133 bp |
| **Contig N90** |  | 77,610 bp |
| **Gaps** |  | 1,113 |
| **Genome BUSCO** | Complete | 95.6% |
|  | Single copy | 92.6% |
|  | Duplicated | 3.0% |
| **Repeat content** |  | 48.06% |
| **LTR** | Gypsy | 0.35% |
|  | unknown | 8.04% |
| **TIR** | CACTA | 1.53% |
|  | Mutator | 20.39% |
|  | PIF Harbinger | 1.37% |
|  | Tc1 Mariner | 0.06% |
|  | hAT | 6.36% |
| **Helitron** |  | 2.10% |
| **Predicted genes** |  | 24,912 |
| **Protein BUSCO** | Complete | 96.3% |
|  | Single copy | 91.9% |
|  | Duplicated | 4.4% |

**Figure 2:**
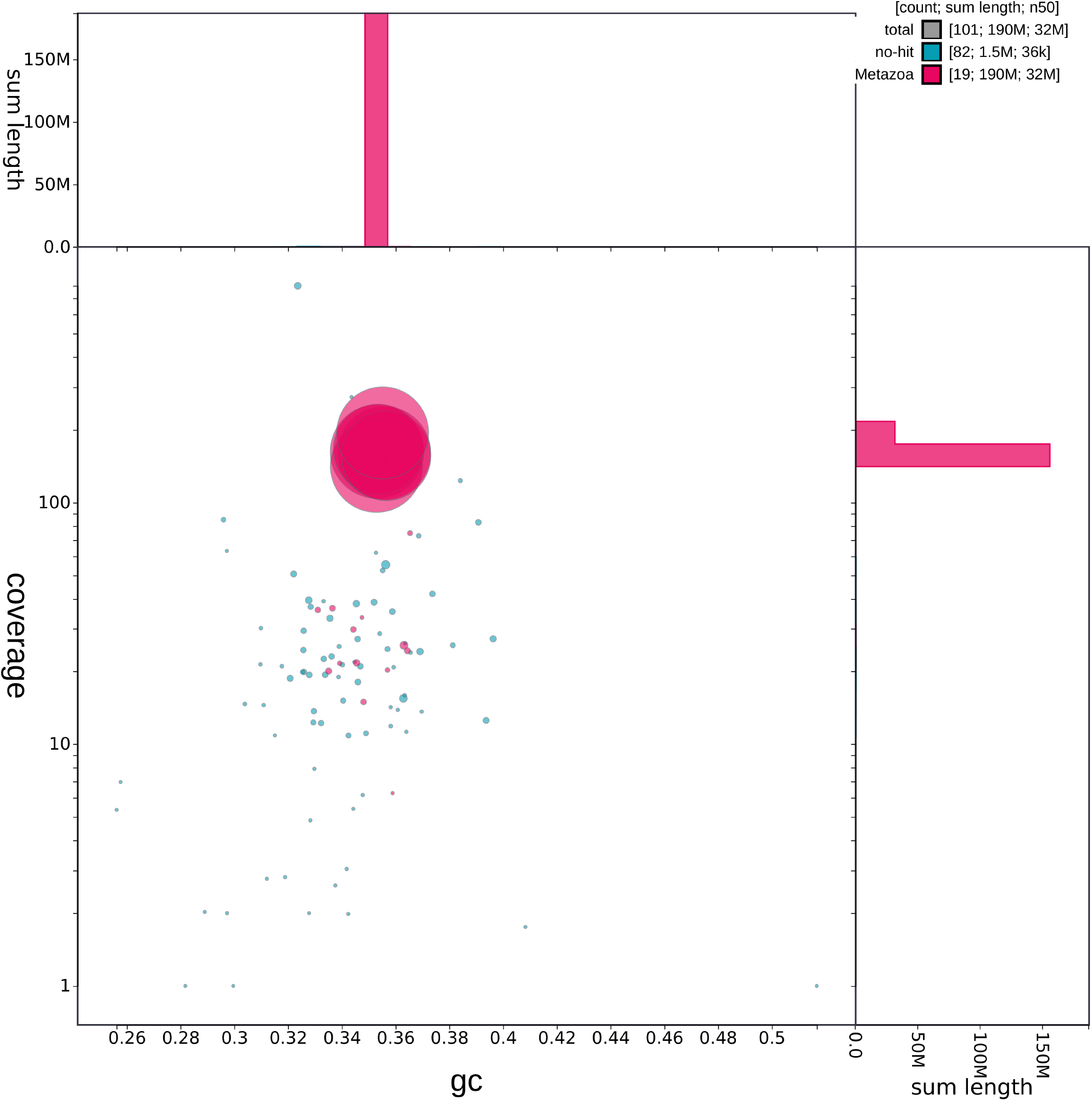
Removal of bacterial contamination from the final assembly. BlobToolKit plot of GC content and PacBio HiFi coverage. Dots are colored following the kingdom to which they were attributed based on BLAST hits (Metazoa in pink, no hit in blue). Bacterial contigs, often present as food source for this species, were removed from the final assembly.

**Figure 3:**
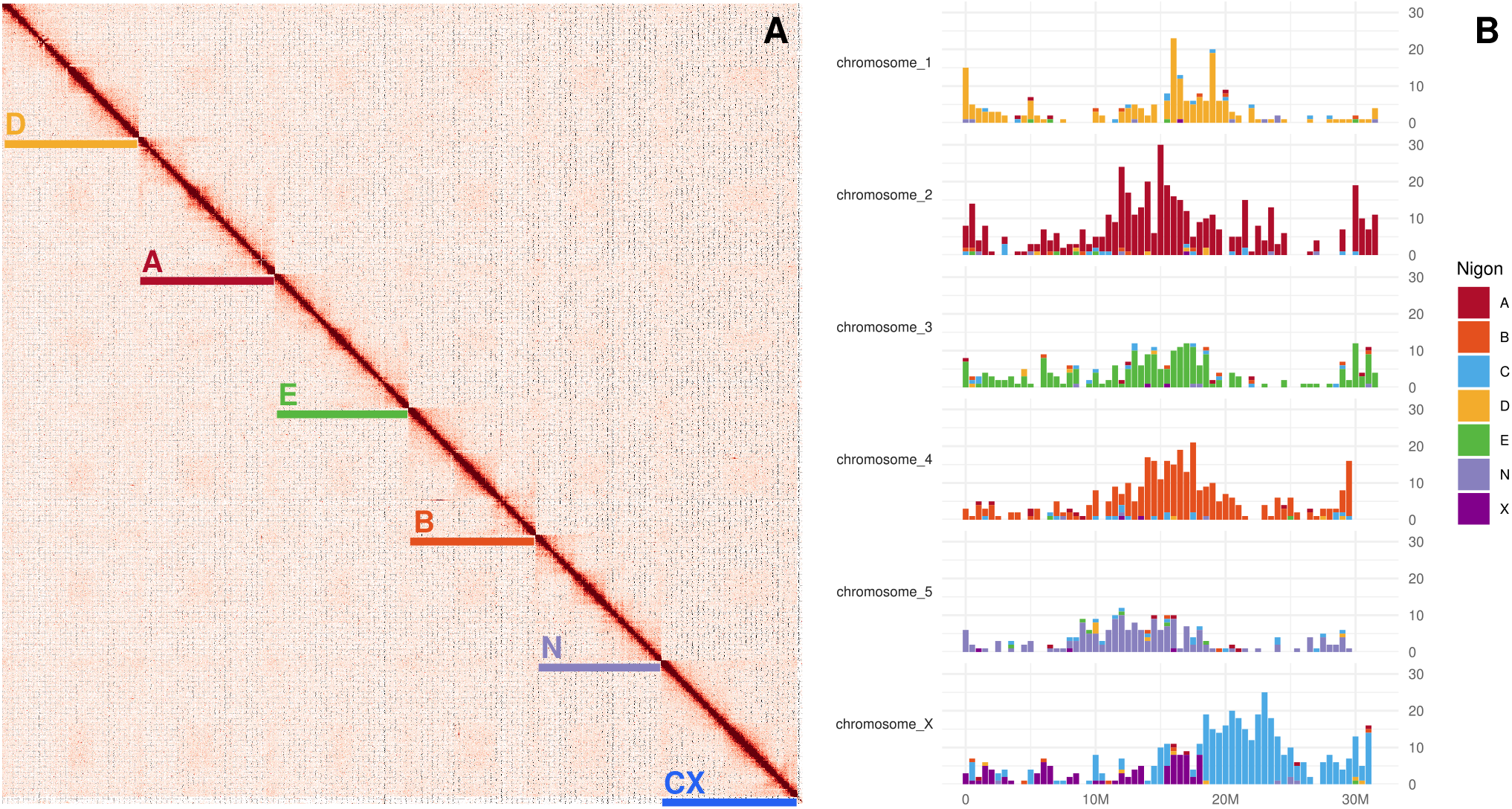
The nuclear genome assembly includes six chromosomes in which Nigon ancestral linkage groups are conserved. A) Hi-C contact map with chromosomes identified based on their Nigon group. B) Identification of Nigon elements for each chromosome.

Comparison of the six chromosomes to the Nigon element model of ancestral chromosomes indicated retention of five of the seven elements as single chromosomes (A, B, D, E, N; Figure 3B). One chromosome comprised a fusion of the C element with the X element (Figure 4). Nigon X in sexual species is the sex chromosome, usually found haploid in males [54, 39]. The assembly was annotated to contain 48.1% repeat. Annotation of transposable elements identified a majority of Mutator-like elements, representing 20.4% of the assembly. 24,912 genes were predicted, with a BUSCO score against the Nematoda odb12 lineage of 96.3% (91.9% single-copy orthologs, 4.4% duplicated orthologs). The mitochondrial genome assembly could not be circularized; it reached 17,634 bp, close to the reference assembly of *Acrobeloides varius* at 17,650 bp.

**Figure 4:**
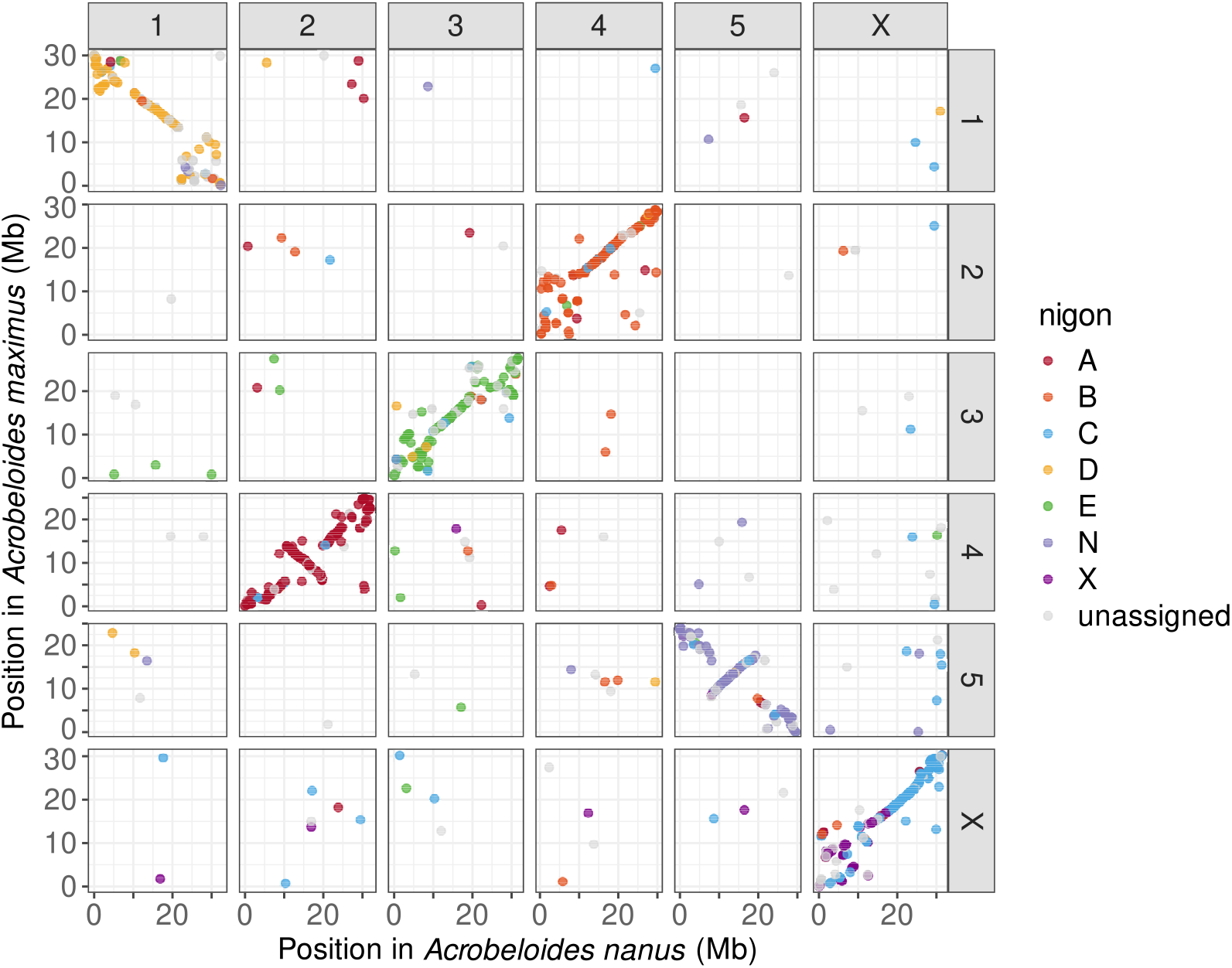
Conservation of six chromosomes in *Acrobeloides nanus* and *Acrobeloides maximus*, with widespread rearrangements. Oxford dot plot of conserved orthologs between the two species, based on the Nigon element model of ancestral linkage groups for nematodes. Dots represent matches between genes and are colored based on their category (A, B, C, D, E, N, X, unassigned).

We compared the chromosomal structure of *A. nanus* with the publicly available assembly of *A. maximus* (GCA 964212105.1), using conserved orthologs from the Nigon element model. Both species have six chromosomes, with one-to-one matches. Some rearrangements are observed, particularly at: the extremities of chromosomes 1 (Nigon D); the extremities of chromosomes 3 (Nigon E); the beginning of chromosome 4 of *A. nanus* and chromosome 2 of *A. maximus* (Nigon B); and the X portion of chromosomes X.

The combination of long reads and Hi-C brought the genome assembly of *Acrobeloides nanus* to chromosome level, far over the kilobase-level contiguity of previous short-read assemblies. Preliminary comparison with *Acrobeloides maximus* revealed an overall conservation of chromosomal organization between the two species, nuanced by multiple intra- and interchromosomal rearrangements. The availability of chromosome-level assemblies for *A. nanus, A. maximus* and more representatives of the family Cephalobidae in the future will bring new insights into their phylogeny, developmental and morphological genes, and the evolution of their genomes.

## Data availability

Sequencing datasets and assembly have been submitted to European Nucleotide Archive under the accession number PRJNA1200840. Annotations are available in the following repository doi.org/10.6084/m9.figshare.31316383.

## Acknowledgements

This project was supported through a DFG Emmy Noether Program (ENP) Projekt (434028868) to PHS and the Excellent Research Support Program of the University of Cologne funding the Biodiversity Genomics Center Cologne (BioC2) led by PHS. NG’s position was first funded through a DFG grant BA 58004-1 to PHS as part of the DFG Sequencing call 2021, and subsequently through the European Union’s Horizon Europe research and innovation programme under the Marie Sklodowska-Curie grant agreement No. 101110569. This work was supported by the DFG Research Infrastructure West German Genome Center (407493903) as part of the Next Generation Sequencing Competence Network (project 423957469). Computational support and infrastructure were provided by the “Centre for Information and Media Technology” (ZIM) at the University of Düsseldorf (Germany). Work at the Wellcome Sanger Institute Tree of Life project is supported by Wellcome Trust award 220540/Z/20/A.

